# Constraints on microbial communities, decomposition and methane production in deep peat deposits

**DOI:** 10.1101/787895

**Authors:** L.A. Kluber, E.R. Johnston, S.A. Allen, J.N. Hendershot, P.J. Hanson, C.W. Schadt

**Affiliations:** Biosciences Division, Oak Ridge National Laboratory, 1 Bethel Valley Rd. Oak Ridge, TN 37831; Climate Change Sciences Institute, Oak Ridge National Laboratory, 1 Bethel Valley Rd. Oak Ridge, TN 37831; Environmental Sciences Division, Oak Ridge National Laboratory, 1 Bethel Valley Rd. Oak Ridge, TN 37831; Department of Microbiology, University of Tennessee, 1311 Cumberland Ave, Knoxville, TN 37996

**Keywords:** Peat, Catotelm, Methane, Microbial communities, Warming, Climate Change

## Abstract

Peatlands play outsized roles in the global carbon cycle. Despite occupying a rather small fraction of the terrestrial biosphere (∼3%), these ecosystems account for roughly one third of the global soil carbon pool. This carbon is largely comprised of undecomposed deposits of plant material (peat) that may be meters thick. The fate of this deep carbon stockpile with ongoing and future climate change is thus of great interest and has large potential to induce positive feedback to climate warming. Recent *in situ* warming of an ombrotrophic peatland indicated that the deep peat microbial communities and decomposition rates were resistant to elevated temperatures. In this experiment, we sought to understand how nutrient and pH limitations may interact with temperature to limit microbial activity and community composition. Anaerobic microcosms of peat collected from 1.5 to 2 meters in depth were incubated at 6°C and 15°C with elevated pH, nitrogen (NH_4_Cl), and/or phosphorus (KH_2_PO_4_) in a full factorial design. The production of CO_2_ and CH_4_ was significantly greater in microcosms incubated at 15°C, although the structure of the microbial community did not differ between the two temperatures. Increasing the pH from ∼3.5 to ∼5.5 altered microbial community structure, however increases in CH_4_ production were non-significant. Contrary to expectations, N and P additions did not increase CO_2_ and CH_4_ production, indicating that nutrient availability was not a primary constraint in microbial decomposition of deep peat. Our findings indicate that temperature is a key factor limiting the decomposition of deep peat, however other factors such as the availability of O_2_ or alternative electron donors and high concentrations of phenolic compounds, may also exert constraints. Continued experimental peat warming studies will be necessary to assess if the deep peat carbon bank is susceptible to increased temperatures over the longer time scales.

## Introduction

Owing to their cool, saturated conditions, northern peatlands serve as an extensive carbon (C) sink storing approximately one third of the world’s terrestrial C (Bridgham et al., 2006; Gorham, 1991; Post et al., 1982). In these systems, peat profiles accumulate largely undecomposed plant material to several meters deep representing thousands of years of C accumulation (Harden et al., 1992; Turunen et al., 2002). Although it has long been established that the peatland C balance is sensitive to anthropogenic disturbance (Armentano & Menges, 1986), there remains considerable uncertainty about how these systems will respond to changes in climate (Melton et al., 2013). Warming trends are expected to be greatest at high latitudes (Collins et al., 2013) and there has been an increased effort to understand how C cycling processes in northern peatlands will respond to these predicted changes. Much of this effort has focused on the acrotelm, the shallow peat that experiences a fluctuating water table, where it is expected that warmer and drier conditions will stimulate C mineralization (Dorrepaal et al., 2009; Gorham, 1991; Ise et al., 2008 Updegraff et al., 2001). The catotelm, saturated and anoxic deep peat that may extend meters below the surface, is not likely to experience the drying that may occur in the shallow peat. However temperature-induced changes in microbial community composition or function in the catotelm layer could dramatically alter the C balance in these systems over time.

As part of the effort to understand ecosystem-level responses to climate changes, the Spruce and Peatland Responses Under Changing Environments experiment (SPRUCE; http://mnspruce.ornl.gov) was designed to achieve whole ecosystem warming of a boreal peatland system (Hanson et al., 2017). Located in the Marcell experimental Forest (Minnesota, USA), this regression-based experiment began with a 13-month deep peat heating treatment, where experimental plots were heated up to +9 °C above ambient conditions from the surface down to a depth of 2 m. Although surface CH_4_ flux was significantly correlated with deep peat heating, results indicated that activity in the surface acrotelm peat, not deep catotelm peat, was responsible for increased CH_4_ production (Wilson et al., 2016). Additionally, Wilson et al. (2016) found that microbial communities and C decomposition did not respond to the 13 months of *in-situ* warming. Findings from this first year of the SPRUCE experiment suggested that the deep peat carbon pool may remain stable despite increased temperature. Ambient temperatures in the deep peat remain relatively stable throughout the year, averaging 6-7 °C between 1.5 to 2 m at the SPRUCE site. The finding that microbial communities in the deep peat did not respond to even the highest (+9 °C) treatment was somewhat surprising given that numerous studies have shown a shift in community structure in response to elevated temperature (DeAngelis et al., 2015; Deslippe et al., 2012; Karhu et al., 2014; Zogg et al., 1997). The structure of peat microbial communities not only determines the functional mechanisms responsible for C decomposition (Cleveland et al., 2014), but may also influence temperature sensitivity of respiration rates (Karhu et al., 2014). Understanding what factors may lead to, or limit, shifts in microbial community structure should aid in constraining C balance of the system and in forming predictions as to the trajectory of these systems under a warming climate.

With low availability of electron acceptors, fermentation and methanogenesis are the primary C decomposition processes in the anoxic catotelm. These processes lead to the accumulation of organic acids and a correspondingly low pH. Ombotrophic bogs typically have a pH ≤ 4.5, well below the reported optimum pH of 5.5-6 for methanogenesis (Kotsyurbenko et al., 2004; Williams & Crawford, 1984). Consequently, acidic conditions force physiological constraints on microbial communities and their activities, such as CH_4_ production. Rain-fed ombotrophic bogs are also characteristically low in nutrients and previous work has indicated that microbial communities can be limited by low nitrogen (N) and phosphorus (P) availability in these systems. Bragazza et al. (2006) saw that *Sphagnum* decompositions rates were correlated with atmospheric N deposition across a European gradient of N deposition, and Tfaily et al. (2014) found that N is immobilized in the highly humified deep peat in the same Minnesota peatland under study here. Using a combination of NMR, enzymatic assays, and metagenome sequencing, other previous work has also found evidence suggesting N and P limitation (Lin et al., 2014a; Steinweg et al., 2018). Hence, nutrient and pH limitations may serve to protect the C bank from elevated temperatures. Conversely, additional perturbations to peatland nutrient cycles, such as N deposition, could have the potential to alter this C balance in peatland systems. Given these uncertainties, the objective of our study was to identify factors that may be limiting decomposition and microbial community change in deep peat deposits. We conducted a 70-day microcosm incubation of deep peat (originating from 150-200 cm depth) at 6 and 15 °C to mimic ambient and +9 **°**C SPRUCE project experimental conditions, with added treatments including elevated pH and the addition of N and P in a full factorial design. Microcosms were monitored for CO_2_ and CH_4_ production, and microbial community dynamics were assessed using qPCR and amplicon sequencing of the bacterial, archaeal, and methanogen communities. We expected that alleviation of each of these potential constraints (pH, N, and P) would lead to increased decomposition as monitored by CO_2_ and CH_4_ production, as well as shifts in microbial community abundance and composition.

## Methods

### Site description and sampling methods

The SPRUCE experiment field site (http://mnspruce.ornl.gov) is located at the Marcell Experimental Forest S1-Bog (N 47°30.4760’; W93°27.162’) in northern Minnesota, USA. Soil in this 8.1 ha ombotrophic peatland has been characterized as the Greenwood series, a Dysic, frigid Typic Haplohemist, (http://www.websoilsurvey.nrcs.usda.gov) with an average depth of 2 to 3 m and pH in the range of 3 to 4.5 that varies with depth. The overstory vegetation consists primarily of *Picea mariana* (Mill.) Britton, Sterns & Poggenburg (black spruce) and *Larix laricina* (Du Roi) Koch (larch) while the understory is dominated by ericaceous shrubs and intermittent herbaceous species with a sphagnum surface layer. The S1-Bog and the SPRUCE experiment has been extensively studied with complete above and belowground characteristics described in previous publications and data products (see Griffiths et al., 2017; Hanson et al., 2016a; Kolka et al., 2011; Tfaily et al., 2014; Wilson et al., 2016). The SPRUCE experiment includes three raised metal boardwalks extending 88, 112, and 92 m into the S1-Bog. The ends of these transects are ∼80 m apart and served as non-treatment areas to sample the deep peat used in this experiment. In August 2016 two cores were taken from each of these three transects with a Russian peat borer (Eijkelkamp, Netherlands). Deep peat (150-200 cm below surface) was homogenized, distributed into one liter Nalgene bottles, topped off with porewater taken from the same depth to a zero headspace, then capped and stored at 4 °C in a coldroom until laboratory microcosm construction.

### Microcosm construction

To examine whether the temperature response of deep peat communities was limited by other environmental factors, a full factorial experimental design with elevated temperature, N, P, and pH treatments was employed with samples from each transect serving as replicates (n=3 for each condition and time point; see below). Each treatment consisted of two levels, control (approximately ambient) and elevated. At our study depth of 150-200 cm, there is low spatial and temporal variation in peat properties (Griffiths et al., 2017) with N, P and pH values averaging 24 g N and 650 mg P per kg dry peat, and a pH of ∼4.5 (Griffiths et al., 2017; Hill et al., 2014; Iverson et al., 2014; Tfaily et al., 2014). Because we wanted to eliminate potential N or P limitations with minimal secondary effects on respiratory processes such as denitrification, 2.54 M NH_4_Cl was used to increase available N by 24 mg N per g dry peat and 29 mM KH_2_PO_4_ was used to increase P availably by 0.65 mg P per g dry peat, effectively doubling N and P without adding potential electron acceptors. Peat pH was adjusted to 5.5 using 0.5 M NaOH for the elevated pH treatment in an effort to alleviate potential pH constraints on CH_4_ production (Ye et al., 2012). Initial 10 day single variable trials were conducted to ensure that nutrient additions did not influence pH and that pH adjustments were stable over time. pH was also verified to be within 0.2 units of target pH on samples taken at 35 & 70 day destructive samplings (data not shown). Homogenization, nutrient additions, and microcosm setup were conducted in a glovebag under anoxic conditions and microcosms were constructed with 5 grams of peat distributed into 120 ml serum vials with butyl rubber stoppers. In total, 96 microcosms were constructed to account for the 16 treatment combinations (N x P x pH x temperature), 2 destructively sampled time points (35 and 70 days), and 3 replicates for each time point. Although all peat samples were saturated at the time of sampling, sterile Milli-Q water was used to equalize the water content after processing to ensure equal water content across sample vials representing the treatments and transect replicates. Temperature treatments were 6 °C, to mimic the SPRUCE ambient plot temperatures, and 15 °C to mimic the SPRUCE +9 °C treatment, with Precision Refrigerated Incubators (Thermo Scientific Inc.). At this depth, *in-situ* peat is largely buffered from temporal temperature fluctuations and remains near these target temperatures year-round (Hanson et al., 2016a; Griffiths et al., 2017). Microcosms were placed inside styrofoam coolers within incubators to reduce the chance of temperature and incubators were maintained within 0.5 °C of target temperatures.

### Quantification of CO_2_ and CH_4_ production

Production of CO_2_ and CH_4_ was determined by sampling headspace gas in three replicate microcosms on days 10, 20, 35, 52, and 70. To avoid introducing biases associated with differing sample sizes, on days 10, 20, and 35, these three replicates for gas sampling were chosen at random (from the six available microcosms), and then on days 52 and 70were done on the three remaining microcosms after destructive harvest for DNA analyses (below). Gas sample measurements were conducted using a SRI 8610C gas chromatograph with a methanizer and flame ionization detector (SRI Instruments) using methods detailed in (Roy Chowdhury et al., 2015). Total CO_2_ and CH_4_ production was calculated as the combined headspace and dissolved gas concentrations (assuming equilibrium) and are reported as μg CO_2_-C or CH_4_-C per g dry peat.

### DNA extraction, quantitative PCR, and rRNA gene amplicon sequencing

Total DNA was extracted from untreated peat samples at the time of setup (time zero) and on samples from destructively sampled microcosms at 35 and 70 days. MoBio PowerSoil DNA extraction kits (formerly MoBio, now Qiagen) were used with a slightly modified protocol that included a 30 minute incubation at 65 °C immediately after the bead beating step. Duplicate 0.25 g samples were extracted and combined during an additional cleanup step using the MoBio PowerClean Pro cleanup kit (MoBio). Cleaned DNA extracts were quantified using a NanoDrop ND-1000 (Thermo Scientific).

Abundance of bacterial, archaeal, and methanogen community members was determined by quantitative PCR (qPCR) of bacterial and archaeal 16S rRNA genes, and methyl coenzyme M reductase (mcrA) genes using previously described methods (Wilson et al., 2016). Bacterial 16S was targeted using Eub338 and Eub518 primers (Lane, 1991; Muyzer et al., 1993), Archaeal 16S was targeted with 915F and 1059R primers (Yu et al., 2005), and mcrA_F and mcrA_R (Luton et al., 2002) were used to target the methanogen specific Methyl–coenzyme M reductase (mcrA) functional gene. Briefly, triplicate reactions were performed on a CFX96TM Real-Time PCR Detection System (Bio-Rad Laboratories) with iQ SYBR Green Supermix (Bio-Rad, CA, USA). Ct values were then compared to standard curves for each gene constructed from *Escherichia coli* and *Methanococcus maripaludis* S2 genomic DNA, corrected for copy number and genome size, and are reported as gene copies per gram of dry peat.

Sequencing of 16S rRNA genes followed the two-step approach described previously (Lundberg et al., 2013) using frameshifting nucleotide primers and barcode tag templates. However, we used a modified primer mixture to increase phylogenetic coverage of bacteria and archaea. The primer mixture consisted of 10 forward and 7 reverse 515F and 806R primer variants (see Cregger et al., 2018) that were combined in equal concentration at 0.5 µM to amplify the V4 region of 16S rRNA gene (Table S1). A 0.7 to 1 ratio of Agencourt AMPure XP beads (Beckman Coulter) were used to clean PCR products prior to the secondary PCR with barcoded reverse primers. Experimental units were then pooled based on agarose gel band intensity to make several replicate libraries and again purified with Agencourt AMPure XP beads. Library quality was then analyzed using a 2100 Agilent Bioanalyzer (Agilent Technologies) and the library with the highest quality was selected for sequencing. After quantification with Qubit (Invitrogen), the amplicon library was adjusted to a final concentration of 9 pM, spiked with 15% PhiX and paired-end sequenced using an Illumina MiSeq (Illumina, San Diego, CA, USA) with 500 (2 x 250) cycles. All amplicon sequence data are available from the National Center for Biotechnology Information under project ID PRJNA362203.

### Data analysis

Data were checked for outliers and normality prior to statistical analysis using R v.3.2.4 (R Core Team, 2016). The qPCR data required square root transformation while all other data were analyzed without transformation. Three-way ANOVAs blocked by sampling site (transect) with Tukey HSD mean comparisons were used to test the effect of temperature, nutrient additions, and pH treatments on alpha diversity, taxa abundance, qPCR gene copies, and cumulative gas production after 70 days. Because the aim of this study was to examine which factors have the ability to influence the structure and function of deep peat microbial communities, results were focused primarily on comparing incubation treatments against the control incubations (6 °C, no addition, ambient pH). All figures were constructed using R package ggplot2 (Wickham, 2009).

Sequence data were processed using a combination of UPARSE and QIIME (v 1.9.1) pipelines (Caporaso et al., 2010; Edgar, 2013). Paired-end sequences were joined, demultiplexed, and screened for quality using the split_libraries_fastq.py command with stringent quality standards (p = 0.95, r = 1, q = 19). Forward and reverse primers were trimmed using Cutadapt (Martin, 2011). The UPARSE pipeline was then used to remove low quality sequences and chimeras prior to 97% OTU clustering with the pick_open_reference_otus.py command that performs closed-reference OTU picking prior to de novo OTU picking. Taxonomy was assigned using the BLAST algorithm against the Greengenes (v 13.9) database (McDonald et al., 2012a; Werner et al., 2012). The resulting OTU table was converted into BIOM format (McDonald et al., 2012b) and singleton and contaminants (mitochondria and chloroplasts) OTUs were removed. To better assess the starting communities, the three time zero samples were sequenced to a greater sequence depth than the 96 incubated samples, ranging from 2-3 million sequences per sample compared to 6 – 60 thousand sequences). Because of this, the time zero samples were separated from the incubation samples prior to threshold filtering and community analysis. Bokulich threshold filtering was employed at 0.005% across the OTU table (Bokulich et al., 2013) and samples were rarefied to 10,000 sequences and represented 687 OTUs. Representative sequences from each OTU were aligned using PyNAST (Caparaso et al., 2010) and a phylogenetic tree was constructed from aligned sequences using FastTree (Price et al., 2009). Alpha and beta diversity metrics were calculated using the alpha_diversity.py and core_diversity.py scripts in QIIME. Subsequent plotting and statistical analyses of alpha-diversity metrics were carried out in R v.3.2.4 (R Core Team, 2016) using the previously described blocked, three-way ANOVA. The R package Vegan v2.3-5 (Oksanen et al., 2016) was used to construct principal coordinate analysis (PCoA) plots with the weighted and unweighted UniFrac distance matrices (Lozupone & Knight, 2005) and to test for community responses to treatments using PERMANOVA analyses with Bray-Curtis distance measures. There were no significant differences between 35 and 70 day harvests in microbial community structure, therefore for the results are only discussed for 70 day samples (see supplemental figures S2-S4 for 35 day data).

## Results

### CO_2_ and CH_4_ production activity

The cumulative amount of CO_2_-C released after 70 days was significantly greater in microcosms incubated at 15 °C (p < 0.001) although neither nutrient additions or pH (p > 0.1) influenced the amount of C released as CO_2_ (Fig. 1). The cumulative amount of CH_4_-C released after 70-days was also greater in the 15 °C incubations (p < 0.001). Although CH_4_-C released was not influenced by pH (p = 0.518), there was a significant effect of nutrient addition (p < 0.001) that was driven by a marked decrease in CH_4_-C production in the treatments with nitrogen addition (N & NP)(p < 0.001 for both addition treatments). The ratio of CH_4_-C to CO_2_-C also increased with elevated temperature (p = 0.016) but decreased with N and NP additions (p < 0.001 for both). There were no significant interaction effects in either the CO_2_ or CH_4_ analysis.

**Figure 1.**
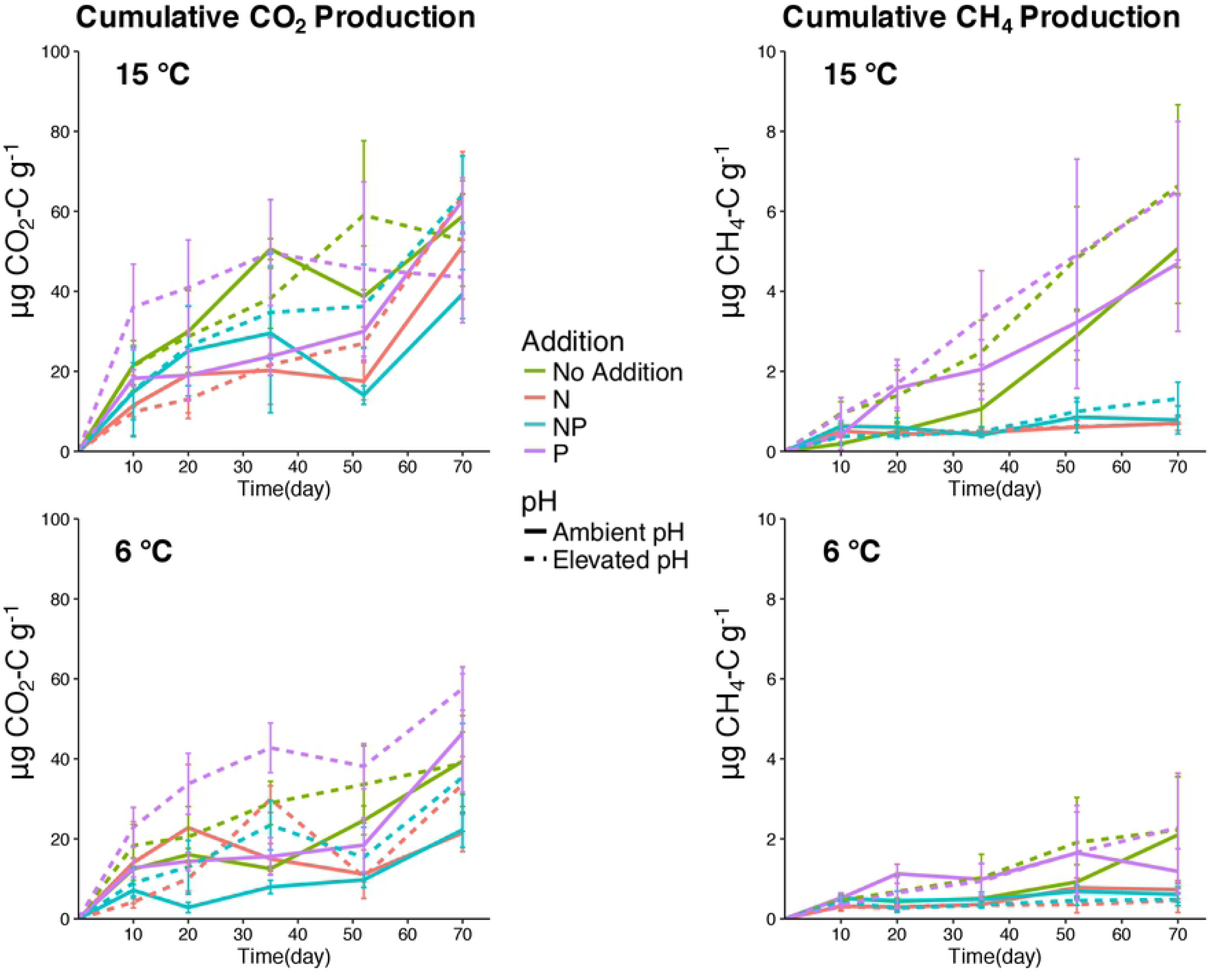
Cumulative production of CO_2_ (left) and CH_4_ (right) in peat microcosms incubated at at 6 (bottom) and 15 °C (top). Nutrient addition treatments are coded by color and the elevated pH treatment is represented by dashed lines.

### Microbial abundance measures with qPCR

Fig. 2 shows the overall response of microbial community members to incubation temperature, nutrient additions, and pH treatments. Data were log transformed prior to analysis with a blocked ANOVA to account for the variation in abundances among the original transect samples that served as replicates in our experiment. Bacterial 16S rRNA gene abundance was not influenced by incubation temperature (p = 0.181) but increased under elevated pH conditions (p < 0.001). Compared to controls, bacterial abundance decreased with N and NP additions under ambient pH (p < 0.001 for both additions). However, under elevated pH conditions, bacterial abundance in N and NP additions were not significantly different from control incubations (p = 0.999 for both). Under ambient pH conditions archaeal 16S rRNA gene abundance decreased with elevated incubation temperature (p = 0.011). However at 15 °C, elevated pH incubations had significantly increased archaeal abundance compared to unamended controls (p = 0.009). The archaeal response to nutrient additions mirrored that of the Bacteria in that incubations receiving N (and NP) addition had significantly lower copy numbers compared to controls (p < 0.001 for both N and NP), while N and NP additions combined with elevated pH were not significantly different from control (p = 0.891 and p = 1.000). Methanogen abundance, measured by quantifying the *mcrA* gene copy number, followed a similar pattern as the archaeal 16S gene abundance. Elevated incubation temperature led to a lower abundance of methanogens under ambient pH conditions (p = 0.001) while methanogen abundance was greater than controls under elevated pH conditions (p < 0.001 for both temperatures). Decreased methanogen abundance was also seen with N and NP additions at ambient pH (p < 0.001 for both additions), but this effect was not evident under elevated pH conditions (p = 0.482 and p = 0.599). Although both bacterial and archaeal abundances decreased with N and NP additions (see above), the ratio of Archaea to Bacteria increased only with N addition (p = 0.038). Elevated incubation temperature reduced the ratio of Archaea to Bacteria (p = 0.001) suggesting that higher incubation temperatures favor bacterial populations; however as mentioned above, the overall bacterial abundance increases with temperature were not significant (p = 0.181). The addition of P alone did not significantly influence the abundance of bacterial, archaeal or methanogen community members.

**Figure 2.**
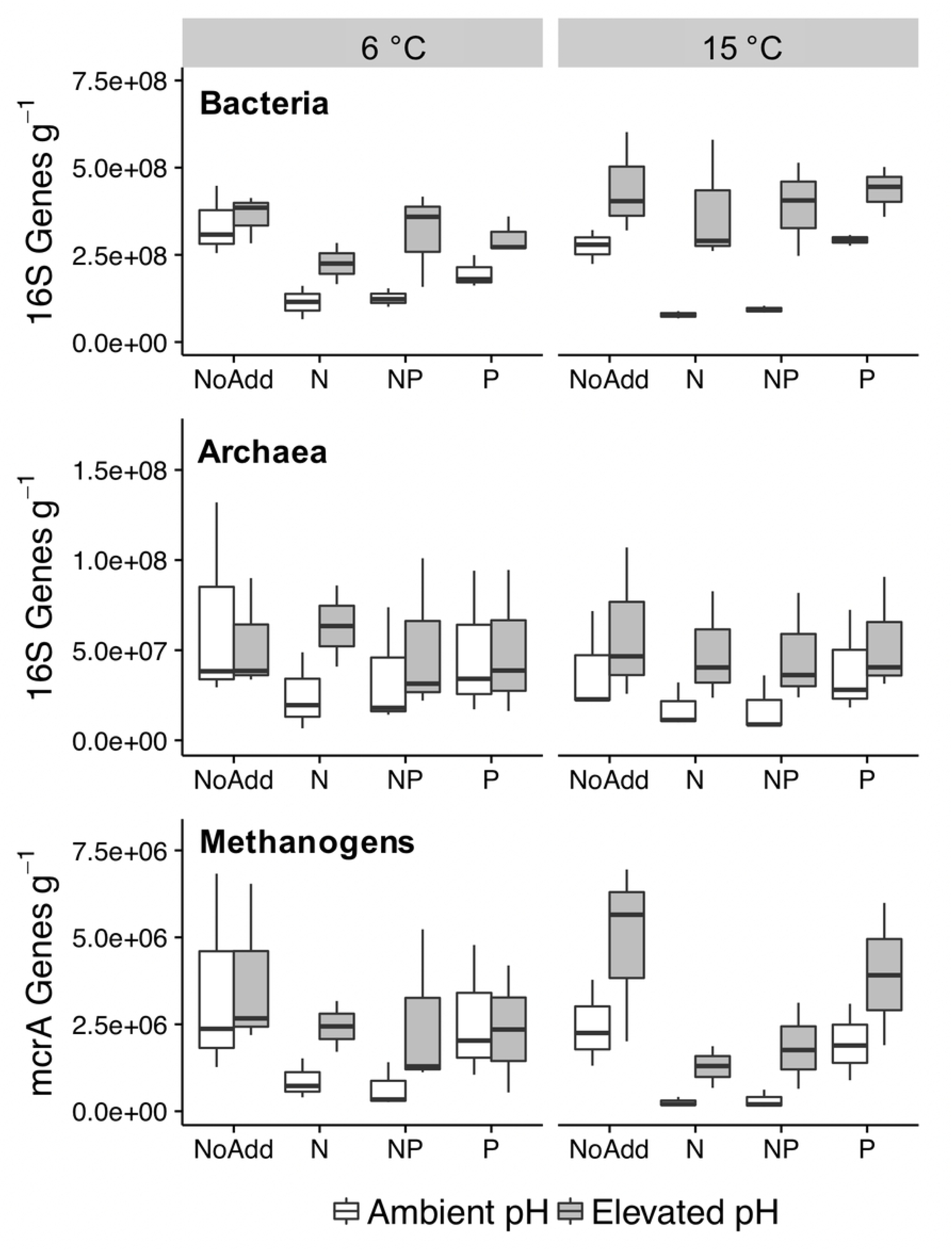
Bacterial, Archaeal, and methanogen abundance measured by qPCR after 70 days of microcosm incubations. Bacterial and archaeal population sizes were determined by quantifying 16S rRNA gene copies and methanogens were assessed by quantifying mcrA gene copies. pH adjustment treatments are shown in grey, and unadjusted treatments are shown in white. Abundance is presented as gene copies per g dry peat.

### Microbial community composition with rRNA-gene sequencing

Both weighted and unweighted PCoA ordinations (Fig. 3) show significant clustering by transect, which was confirmed by PERMANOVA analysis (p < 0.001 for both). Despite the distinct transect clustering pattern within the unweighted ordination, pH also had a significant influence on community composition in both the weighted (p = 0.001) and unweighted (p = 0.002) ordinations and tends to separate along PC1. Driven by the N and NP additions, nutrient addition was a significant grouping factor in the weighted ordination (p = 0.001), although there was not a significant effect of nutrient addition in the unweighted ordinations. Both ordinations and PERMANOVA tests indicated that the community structure was not influenced by temperature or incubation time. After 70 days of incubation, microcosm alpha diversity metrics (Chao1 and Observed OTUs) were significantly lower in the elevated pH treatments (p < 0.001 for both) but did not vary significantly among nutrient addition treatments (Fig. 4). There was also significant spatial variation in alpha diversity metrics (Chao1 p = 0.022 and observed OTUs p = 0.001). Microcosm replicates with peat originally from transect 1 had significantly greater diversity than the other two transects, even after 70 days of incubation. This trend is also seen in peat samples that were harvested at time zero, although only one sample from each transect was sequenced (Fig. S2). Similar results were seen at 35 days of incubation (Fig. S3) with diversity in transect 1 being higher than the other transects and elevated pH treatments having lower diversity than ambient pH. Similar to the community composition measures, these alpha-diversity metrics did not change with incubation temperature.

**Figure 3.**
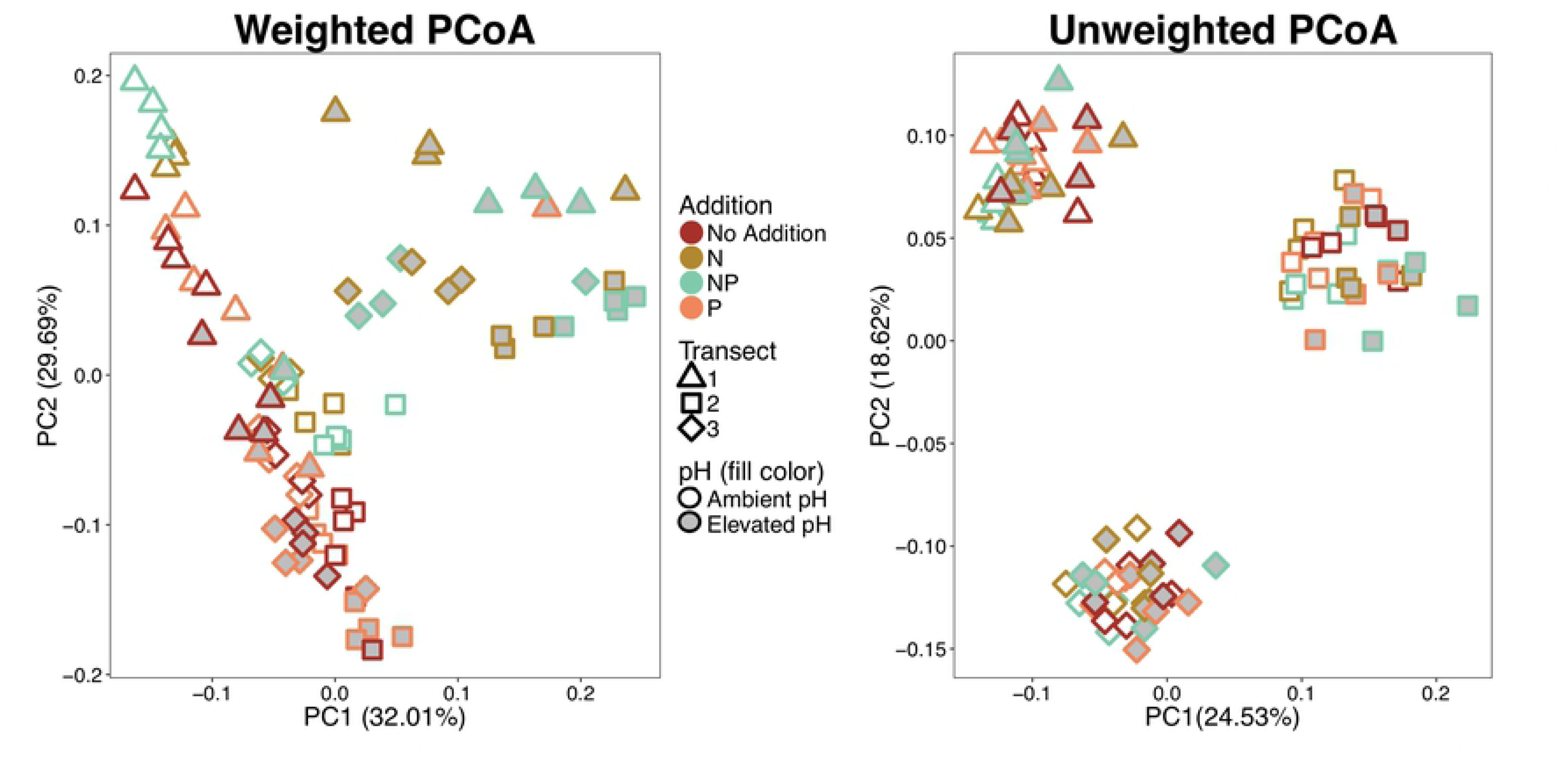
Weighted and unweighted UniFrac PCoA ordinations of microcosm microbial communities. Each point represents an individual microcosm. Color, shape, and fill are used to code for by nutrient addition, transect (origin location), and pH treatment, respectively. Ordinations include samples from both temperature treatments and time points (not denoted) as they did not significantly differ.

**Figure 4.**
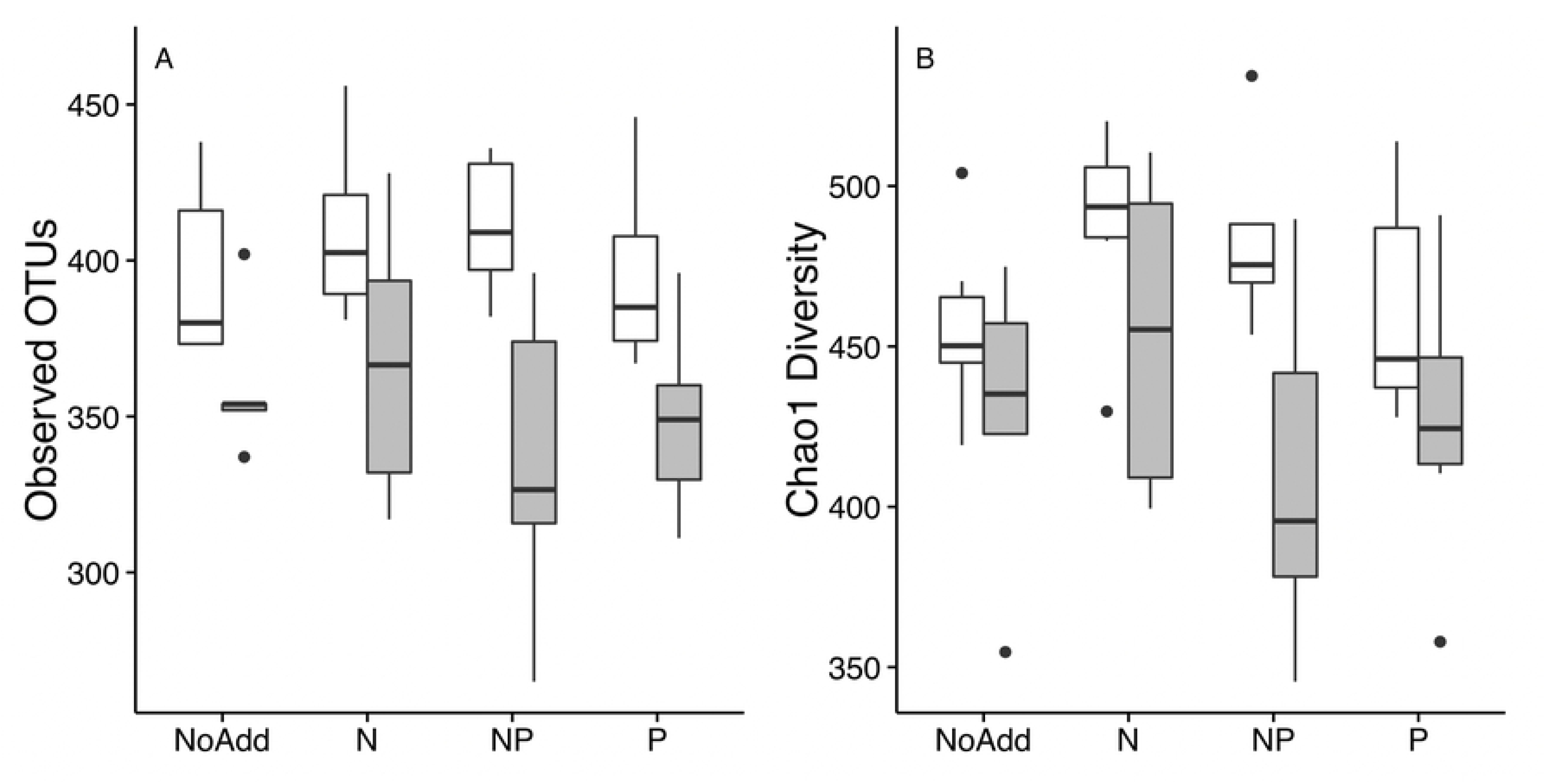
Alpha diversity metrics of microcosm communities after 70 day incubations. pH adjustment treatments are shown in grey, while unadjusted treatments are shown in white. Treatments incubated at 6 and 15 °C were not significantly different and are combined for simplicity, thus each bar represents 6 microcosms.

Relative abundance of dominant phyla and Proteobacteria classes present at the end of the incubation period were primarily influenced by the interactive effects of nutrient addition and pH treatment (Table 1). The significant differences occur only with N or NP additions combined with Elevated pH, where the treatments a had a significantly lower relative abundance of Acidobacteria and Bacteroidetes that corresponds with a substantial increase in the relative abundance in Gammaproteobacteria. This is congruent with the community analysis where the N addition and Elevated pH samples were significantly clustered and divergent from other samples (Fig. 3). Incubation temperature had little effect on the relative abundance of dominant taxa. In fact, the only phyla with a significant main-effect temperature response was Bacteroidetes, where mean relative abundance was slightly, but significantly, greater in samples incubated at 6 °C, rather than samples at 15 °C (6.87% vs. 5.64%, respectively; p = 0.042). Other than the significant interactions discussed above and presented in Table 1, no taxa were significantly influenced by nutrient addition alone. However, elevating pH had a significant negative effect on several taxa; including Chloroflexi, Fibrobacteres, Alphaproteobacteria, and Verrucomicrobia that all had lower relative abundances in incubations with elevated pH than incubations carried out at ambient pH (p-values < 0.001; Table S2).

**Table 1:**
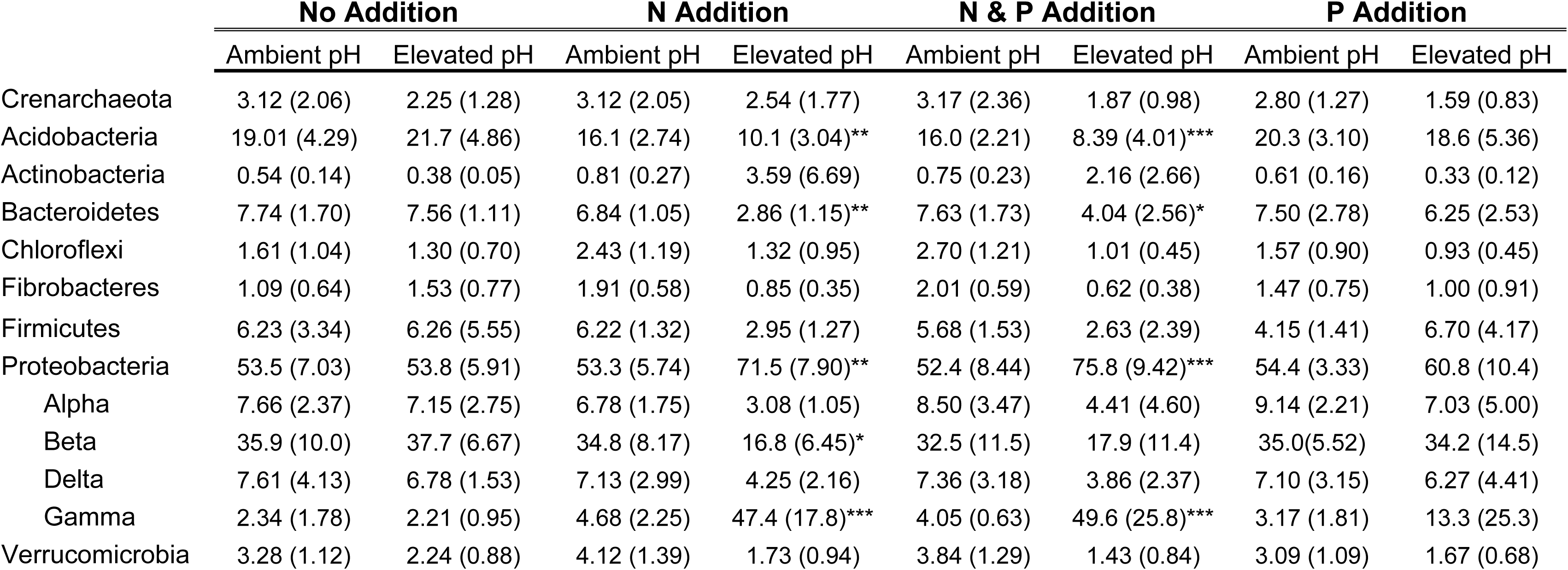
Mean percent relative abundance (and standard deviation) of dominant microbial phyla and Proteobacteria classes after 70 day anaerobic incubation. Data represent combined values for 6 and 15 °C temperature treatments as temperature had little effect on overall microbial community composition and Bacteroidetes was the only taxa to have significant temperature response (p=0.04). Asterisks denote significant difference from the control samples (no addition, ambient pH) determined by Tukey adjusted p-values denoted as ***, **, and * for p = 0.001, 0.01, and 0.05, respectively.

|  | No Addition |  | N Addition |  | N & P Addition |  | P Addition |  |
| --- | --- | --- | --- | --- | --- | --- | --- | --- |
|  | Ambient pH | Elevated pH | Ambient pH | Elevated pH | Ambient pH | Elevated pH | Ambient pH | Elevated pH |
| Crenarchaeota | 3.12 (2.06) | 2.25 (1.28) | 3.12 (2.05) | 2.54 (1.77) | 3.17 (2.36) | 1.87 (0.98) | 2.80 (1.27) | 1.59 (0.83) |
| Acidobacteria | 19.01 (4.29) | 21.7 (4.86) | 16.1 (2.74) | 10.1 (3.04)** | 16.0 (2.21) | 8.39 (4.01)*** | 20.3 (3.10) | 18.6 (5.36) |
| Actinobacteria | 0.54 (0.14) | 0.38 (0.05) | 0.81 (0.27) | 3.59 (6.69) | 0.75 (0.23) | 2.16 (2.66) | 0.61 (0.16) | 0.33 (0.12) |
| Bacteroidetes | 7.74 (1.70) | 7.56 (1.11) | 6.84 (1.05) | 2.86 (1.15)** | 7.63 (1.73) | 4.04 (2.56)* | 7.50 (2.78) | 6.25 (2.53) |
| Chloroflexi | 1.61 (1.04) | 1.30 (0.70) | 2.43 (1.19) | 1.32 (0.95) | 2.70 (1.21) | 1.01 (0.45) | 1.57 (0.90) | 0.93 (0.45) |
| Fibrobacteres | 1.09 (0.64) | 1.53 (0.77) | 1.91 (0.58) | 0.85 (0.35) | 2.01 (0.59) | 0.62 (0.38) | 1.47 (0.75) | 1.00 (0.91) |
| Firmicutes | 6.23 (3.34) | 6.26 (5.55) | 6.22 (1.32) | 2.95 (1.27) | 5.68 (1.53) | 2.63 (2.39) | 4.15 (1.41) | 6.70 (4.17) |
| Proteobacteria | 53.5 (7.03) | 53.8 (5.91) | 53.3 (5.74) | 71.5 (7.90)** | 52.4 (8.44) | 75.8 (9.42)*** | 54.4 (3.33) | 60.8 (10.4) |
| Alpha | 7.66 (2.37) | 7.15 (2.75) | 6.78 (1.75) | 3.08 (1.05) | 8.50 (3.47) | 4.41 (4.60) | 9.14 (2.21) | 7.03 (5.00) |
| Beta | 35.9 (10.0) | 37.7 (6.67) | 34.8 (8.17) | 16.8 (6.45)* | 32.5 (11.5) | 17.9 (11.4) | 35.0(5.52) | 34.2 (14.5) |
| Delta | 7.61 (4.13) | 6.78 (1.53) | 7.13 (2.99) | 4.25 (2.16) | 7.36 (3.18) | 3.86 (2.37) | 7.10 (3.15) | 6.27 (4.41) |
| Gamma | 2.34 (1.78) | 2.21 (0.95) | 4.68 (2.25) | 47.4 (17.8)*** | 4.05 (0.63) | 49.6 (25.8)*** | 3.17 (1.81) | 13.3 (25.3) |
| Verrucomicrobia | 3.28 (1.12) | 2.24 (0.88) | 4.12 (1.39) | 1.73 (0.94) | 3.84 (1.29) | 1.43 (0.84) | 3.09 (1.09) | 1.67 (0.68) |

## Discussion

Peatland ecosystems occupy a rather small fraction of the terrestrial biosphere yet account for roughly one third of global soil carbon pool (Bridgham et al., 2006). The decomposition of this carbon stock under ongoing and future climate change has the potential to induce positive feedbacks to warming. In this laboratory study, we found that the mineralization of deep peat C to CO_2_ and CH_4_ was largely controlled by functional, rather than structural, responses of the microbial community to increased temperature. Elevated temperature increased production of both CO_2_ and CH_4_, as well as the ratio of CH_4_:CO_2_ produced. Despite these physiological responses of increased C mineralization, elevated temperature had a relatively minor influence on the microbial community composition. qPCR analyses showed both overall archaeal and methanogen population sizes decreased with elevated temperature, although the size of the bacterial population showed no response. Similarly, the overall community structure was not significantly influenced by temperature. Contrastingly, elevating pH led to significant shifts in peat community structure, and effects of on CO_2_ and CH_4_ production trended higher, but were not statistically significant. The addition of N reduced CH_4_ production and the size of bacterial and archaeal populations. Although population sizes were less impacted by N addition combined with elevated pH, the structure of the microbial community experienced a dramatic shift and CH_4_ production remained significantly lower than the control microcosms.

### Controls on CO_2_ and CH_4_

The well-established relationship between temperature and soil respiration (Raich & Schlesinger, 1992) holds true in peatland systems with Yavitt et al. (1987) describing temperature as the “master variable” controlling CO_2_ production in *Sphagnum*-derived peat bogs. Numerous studies have shown similar results and while there is little doubt that elevating peat temperature will increase peatland CO_2_ emission, the source of the C remains unclear. Dorrepaal et al. (2009) reported elevated CO_2_ emissions in an in-situ warming experiment and attributed 69% of the additional CO_2_ to the deeper peat layers. However, Wilson et al. (2016) found that only peat from the acrotelm produced more CO_2_ under elevated temperatures. It is estimated that over 75% of the C in the S1 bog ecosystem is held within the deeper catotelm layers below 50cm (Griffiths et al., 2017; Iverson et al., 2014). The competing C mineralization processes of fermentation and methanogenesis further complicate understanding of the C balance in peatland systems. While both CO_2_ and CH_4_ emissions increase with temperature (Treat et al., 2015), the relationship between CH_4_ and temperature is not as strong as for CO_2_, leading Yavitt et al. (1987) to suggest that CH_4_ production was limited by factors other than temperature. Isotopic evidence indicates that acetoclastic methanogenesis dominates in shallow peat while hydrogenotrophic methanogensis dominates in deeper peat (Wilson et al., 2016). Both of these pathways are inhibited under low pH conditions leading to increased CH_4_ production in peat incubations with elevated pH (Williams & Crawford, 1984; Ye et al., 2012). Elevated pH treatments consistently trended toward greater CH_4_ production (Fig 1), however likely due to the large differences between our replicates from that were taken from different parts of the bog, these were not significant. This agrees with the suggestion by Ye et al. (2012) that factors other than pH can be responsible for the relatively low CH_4_ production in ombotrophic peatlands, compared to more minerotrophic systems. However, while both Williams and Crawford (1984) and Ye et al. (2012) observed significant increases in methanogenesis above pH 5, they also found peak production to be at pH 6.0 and 6.5 respectively, suggesting that further increases may be needed to better alleviate this constraint.

Although N limitation has been implicated in suppressing C decomposition in ombotrophic peatlands, these studies are primarily focused on surface processes and the evidence is far from conclusive. Nutrient addition and gradient studies have shown that addition of exogenous N increases decomposition and CO_2_ emissions in peat bogs (Bragazza et al., 2006), while others found N additions have no or mixed effects (Aerts & Caluwe, 1999; Aerts et al., 2006; Bubier et al. 2007), or even suppress decomposition (Aerts & de Caluwe 1999; Amador & Jones, 1993; Bubier et al. 2007). The addition of N did not lead to increased CO_2_ or CH_4_ production in our study of deep peat deposits. In fact, N addition had a detrimental effect on the microbial community leading to decreased population sizes, altered community composition, and decreased C mineralization. We intentionally selected NH_4_ as a N source to avoid stimulating denitrification, and known anaerobic ammonium oxidizing organisms were not among those identified as shifting in rRNA-gene based analyses. The mechanism of this effect of N addition remains unresolved by our study, however primarily from the study of other systems such as anaerobic digestors and sewage plants, NH_4_ is known to be an inhibitor of anaerobic processes and methanogenesis at high addition rates (reviewed by Chen et al., 2009; Yenigün & Demirel 2013) and similar phenomena may have contributed to our observations as well. It is possible that the level of N addition used in this study was toxic to some members of the peat microbial community adapted for N limitation, and that a more subtle increase in N would have resulted in a differing response. Molecular and enzymatic data have suggested the possibility of P limitation in the S1 bog (Hill et al., 2014; Lin et al., 2014a, 2014b; Steinweg et al., 2018), and like most ombrotrophic bog ecosystems, peat and porewater samples from the SPRUCE site are low in N and P (Griffiths & Sebestyen, 2016; Griffiths et al., 2017). However, P did not seem to alleviate constraints on CO_2_ or CH_4_ production in our deep peat microcosms at the levels added here, as the addition of P had no significant effect on gas production or microbial community structure. Our findings show that the low nutrient conditions are not likely a primary factor limiting decomposition in the deep peat C bank, thus this large deep peat stock may be subject to accelerated decomposition under the increased temperature of the SPRUCE experiment given further time.

### Role of microbial community

Similar to the findings of Wilson et al. (2016), we did not see overall changes in community composition in response to elevated temperatures, although archaeal, including methanogen, population abundance decreased slightly in qPCR measures. Interestingly, these temperature suppressed methanogen populations were apparently offset by increased metabolic activity resulting in significantly increased CH_4_ production. Organic carbon mineralization to CO_2_ was similarly uncoupled from microbial population levels as measured by qPCR. Other studies have found significant correlations of CH_4_ production to methanogen population activity levels using RNA based analyses (Freitag et al., 2010; Putkinen et al., 2009; Putkinen et al., 2014). Although few studies have examined the responses of microbial communities to peat warming, especially from the deeper layers, many such experiments have been conducted in mineral soils. A meta-analysis of 75 manipulative experiments found that the response of soil communities to warming experiments were best explained by local climate and ecosystem type (Blankinship et al., 2011). Although peat community structure hasn’t responded to ex-situ or in-situ warming, continued long-term warming of SPRUCE plots may lead to altered community structure much like the delayed response seen in other long term soil warming experiments in upland soils (DeAngelis et al., 2015; Melillo et al., 2017; Johnston et al., 2019). However, while peatland microorganisms could be considered ‘slow growers’ and thus need more time for observable shifts in community structure, we did witness significant shifts in community indicators with other experimental factors, suggesting a lag between physiological response and community structure that is specific to increased temperature.

Elevated pH had significant effects on the microbial population sizes, with greater abundances of all microbial groups examined using qPCR. Although pH did influence community structure, the largest treatment effect was seen in microcosms with elevated pH and N addition (elevated pH + N and elevated pH +NP). The combined effect of elevated pH and N addition decreased the abundance of most community members while increasing a single genera, *Rahnella*. Sequences from this a psychrotolerant Gammaproteobacteria comprised 40-50% of the sequence libraries from microcosms with elevated N and pH. Soil pH is known to be an important variable in determining microbial community structure (Fierer & Jackson, 2006; Lauber et al., 2009) and thus a community response to pH was expected. In addition to constraining overall microbial communities, physiological adaptations allowing microbes to survive in this low pH, anoxic environment appear to come at the cost of lower biocatalytic activity compared to neutral pH anoxic sediments (Goodwin & Zeikus, 1987). Microcosms with elevated pH trended toward elevated CO_2_ and CH_4_ production (Fig. 1), however these results were not significant due to high variability between replicates and thus further analysis of its’ limiting effect on the decomposition of deep peat is warranted.

Although methanogen abundance and activity in the bog profile peaks around 30-50 cm in the acretolm (Wilson et al., 2016), our findings show that elevated temperatures have the potential to increase CH_4_ production in the deep peat. Whether CH_4_ produced in the deep peat will diffuse to the surface and be released to the atmosphere, or perhaps be consumed by methane-oxidizing taxa, remains to be determined. Others have reported that *Sphagnum*-associated methane-oxidizers are ubiquitous across in *Sphagnum* bogs and methane-oxidation rates increase with temperature (Freitag et al., 2010; Kip et al., 2010; Putkinen et al., 2014). However the ability of methane oxidation to reduce peatland CH_4_ emissions is largely dependent on both water level and the extent of the temperature increase (van Winden et al., 2012).

The majority of work examining microbial communities in peatlands has focused on *Sphagnum* and plant-associated microbes (Kip et al., 2010; Kostka et al., 2016) and near surface peat (Lin et al., 2012; Preston et al., 2012; Puglisi et al., 2014). Wilson et al. (2016) examined peat communities down to 200 cm from across the SPRUCE experimental site and found that spatial heterogeneity was largely overshadowed by the strong depth stratification. Despite the relatively homogenous physical and chemical properties of deep peat (Iverson et al., 2014; Tfaily et al., 2014), pore water chemistry shows greater spatial variability at depth (Griffiths & Sebestyen, 2016; Griffiths et al., 2017) thus suggesting that low lateral and vertical flow of porewater could limit microbial dispersal and contribute to patchy distributions (Nemergut et al., 2013). Indeed, porewater movement, and thus substrate transport and availability, may be a key difference between our previous in-situ results and this microcosm study. The artificial conditions and homogenization of the peat matrix in our microcosm setup could have increased substrate availability, and may help explain increased CO_2_ and CH_4_ production in the absence of microbial community change. The aim of this study was not to examine microbial communities across the entire bog ecosystem, but rather to examine how peat communities respond to warming and which factors may limit a structural and/or functional response. However, peat from three separate transects was used for microcosm replicates and sequencing revealed distinct communities at each location. Further, the transect blocking variable was significant in the analysis of archaeal and methanogen populations and CH_4_ production. Altogether, these findings suggest that although peatlands will contain sites with differential microbial composition capable of greater CH_4_ production, distinct communities from each transect could have similar functional and structural responses to treatments.

While the surface peat may be most sensitive to elevated temperatures (McKenzie et al., 1998; Williams & Crawford, 1984; Wilson et al., 2016), our findings support Dorrepaal et al. (2009) in showing that the deep C bank is indeed at risk under elevated temperatures. We found that temperature is a factor limiting microbial activity in deep peat reserves. However, additional factors such as low initial population sizes, substrate availability, etc. are likely limiting in-situ decomposition of these deep peat layers, and how these respond to the longer-term SPRUCE manipulations, remains for additional study. Despite a relatively stable microbial community structure and population size, microorganisms inhabiting the deep peat seem to have the potential to become more active under elevated temperatures and thus accelerate the mineralization of the peatland C bank. These results show that elevating temperature can also increase the relative amount of C being mineralized to CH_4_, thus having the potential of further accelerating the feedback loop of climate warming.

## Acknowledgements

The authors would like to thank Taniya Roy Chowhury and Jana Phillips for assistance in designing and implementing the experiment, as well as Zamin Yang, Melissa Cregger, and Allison Veach for assistance with the sequencing and bioinformatic analysis. Further, we thank the USDA Forest Service, Northern Research Station for enabling access to and use of the S1-Bog of the Marcell Experimental Forest. This work was supported by the US Department of Energy, Office of Science, Office of Biological and Environmental Research as part of the SPRUCE project (http://mnspruce.ornl.gov). Oak Ridge National Laboratory is managed by UT-Battelle, LLC, for the U.S. Department of Energy under contract DE-AC05-00OR22725.

